# Antigen-specific stimulation and expansion of CAR-T cells using membrane vesicles as target cell surrogates

**DOI:** 10.1101/2021.03.18.435976

**Authors:** V.M. Ukrainskaya, Y.P. Rubtsov, D.S. Pershin, N.A. Podoplelova, S.S. Terekhov, R.S. Kalinin, I.A. Yaroshevich, A.I. Sokolova, D.V. Bagrov, E.A. Kulakovskaya, V.O. Shipunova, S.M. Deyev, E.G. Maksimov, O.V. Markov, A.L. Oshchepkova, M.A. Zenkova, J. Xie, A.G. Gabibov, M.A. Maschan, A.V. Stepanov, R.A. Lerner.

**Affiliations:** M.M. Shemyakin and Yu.A. Ovchinnikov Institute of Bioorganic Chemistry of the Russian Academy of Sciences, Moscow, 117997 Russia; Dmitry Rogachev National Medical Research Center of Pediatric Hematology, Oncology and Immunology, Moscow, 117997 Russia; Lomonosov Moscow State University, Faculty of Biology, 119991, Moscow, Russia; Federal Research and Clinical Center of Physical-Chemical Medicine of Federal Medical Biological Agency, Russia, 119435, Moscow, Malaya Pirogovskaya, 1a; Institute of Chemical Biology and Fundamental Medicine, Siberian Branch of the Russian Academy of Sciences, Lavrentiev Ave. 8, 630090 Novosibirsk, Russian Federation; Faculty of Biology and Biotechnology, National Research University Higher School of Economics, Moscow, 101000 Russia; Research Centrum for Oncotheranostics, Research School of Chemistry and Applied Biomedical Sciences, Tomsk Polytechnic University, 634050 Tomsk, Russia; Department of Chemistry, The Scripps Research Institute, 10550 North Torrey Pines Road MB-10, La Jolla, CA 92037, USA

**Author notes:** These authors contributed equally to this work (V.M. Ukrainskaya, Y.P. Rubtsov). To whom correspondence should be addressed. Correspondence: Michael M. Maschan, Dmitry Rogachev National Medical Research Center of Pediatric Hematology, Oncology and Immunology, Moscow, 117997 Russia, Alexey V. Stepanov, M.M. Shemyakin and Yu.A. Ovchinnikov Institute of Bioorganic Chemistry of the Russian Academy of Sciences, Moscow, 117997 Russia; Dmitry Rogachev National Medical Research Center of Pediatric Hematology, Oncology and Immunology, Moscow, 117997 Russia; Department of Chemistry, The Scripps Research Institute, 10550 North Torrey Pines Road MB-10, La Jolla, CA 92037, USA, Richard A. Lerner, Department of Chemistry, The Scripps Research Institute, 10550 North Torrey Pines Road MB-10, La Jolla, CA 92037, USA.

## Abstract

Development of CAR-T therapy led to immediate success in the treatment of B cell leukemia and lymphoma. It also raised an opportunity to design new protocols to target solid tumors. Manufacturing of therapy-competent functional CAR-T cells needs robust protocols for *ex vivo/in vitro* expansion of modified T-cells. This step is challenging, especially if non-viral low efficiency delivery protocols are used to generate CAR-T cells. Modern protocols for CAR-T cell expansion are based on incubation with high doses of recombinant cytokines to support proliferation, non-specific stimulation with surface-bound antibodies to induce TCR cross-linking, or co-cultivation with antigen-expressing feeder cell lines. These approaches are imperfect since non-specific stimulation results in rapid outgrowth of CAR-negative T cells, and removal of feeder cells from mixed cultures necessitates additional purification steps. In an effort to develop a specific and improved protocol for CAR-T cell expansion, we took advantage of cell-derived membrane vesicles, and the simple structural demands of the CAR-antigen interaction. Our approach was to make antigenic microcytospheres from common cell lines stably expressing surface-bound CAR antigens (antigenic vesicles, AVs), and then use them for stimulation and expansion of CAR-T cells. We developed a rapid, simple, efficient, and inexpensive protocol to generate, stabilize and purify AVs. As proof-of-concept we tested the efficacy of our AV constructs on several CAR-antigen pairs. The data presented in this article clearly demonstrate that our protocol produced AVs with the capacity to induce stronger stimulation, proliferation and functional activity of CAR-T cells than is possible with existing protocols. We predict that this new methodology will significantly improve the ability to obtain improved populations of functional CAR-T cells for therapy.

**Graphical abstract:** 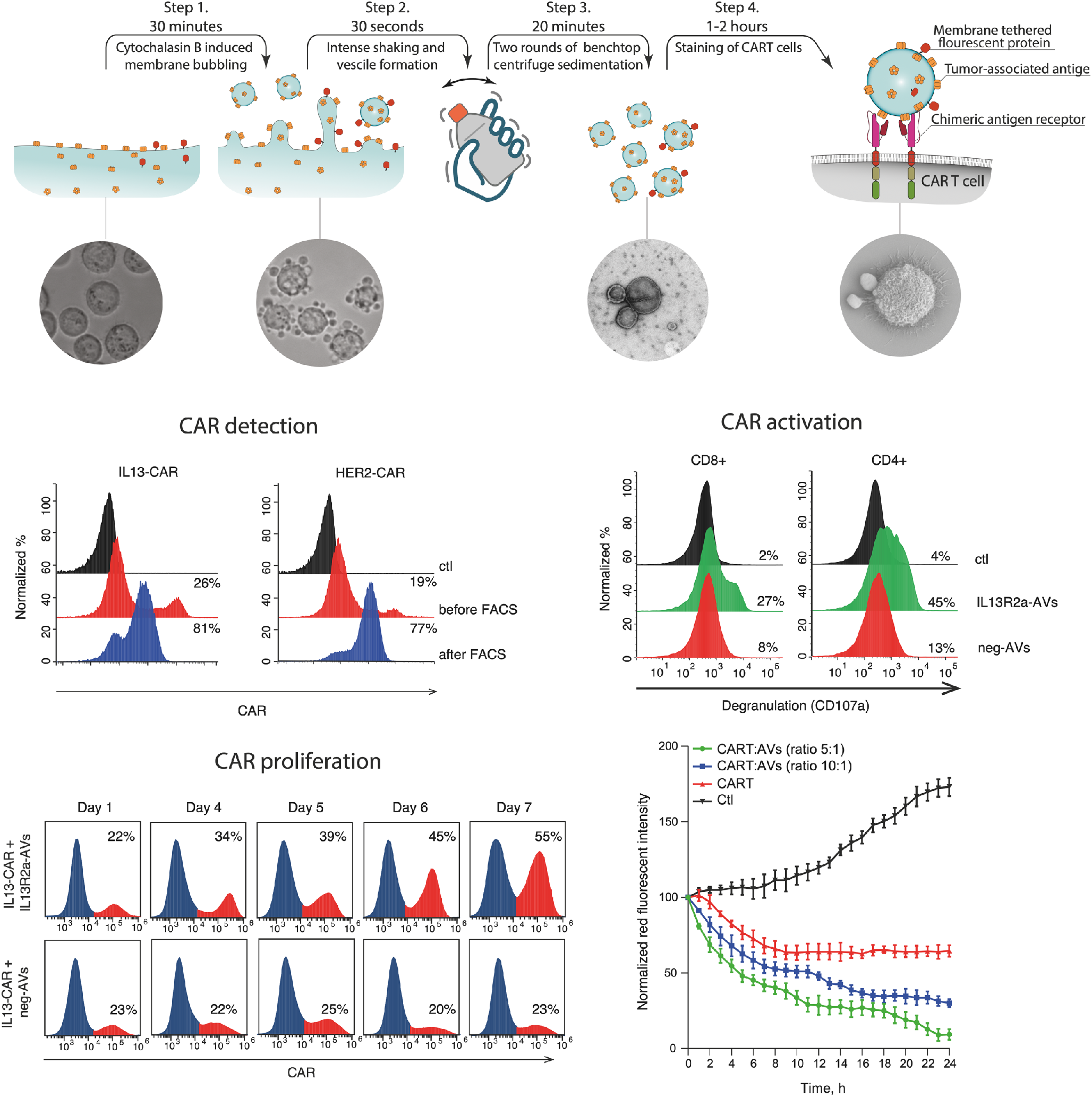

## Introduction

One of the significant challenges in CAR-T therapy is obtaining an adequate number of CAR-T cells with the desired function through donor T cell stimulation and expansion *in vitro*. The majority of existing approaches are based on non-specific or antigen-presenting/feeder cell-based protocols. Non-specific protocols involve the combination of TCR cross-linking with antibodies recognizing constant parts of the TCR and co-stimulatory molecules (anti-CD3 and anti-CD28), with high doses of cytokines, such as IL-2, IL-15 and others^1^. Cross-linking antibodies are usually immobilized on the surface of polymer beads or, in some reports, on sophisticated nanotube scaffolds^2–4^. The main drawback of such non-specific expansion is that CAR-negative ballast cells are also expanding. Thus, if CAR-positive cells are originally present in a low percentage they may be significantly diluted. Another problem is that stimulation of T cells with beads and cytokines results in terminal differentiation of naïve cells into effector T cells with exhausted phenotype^5,6^. Antigen-specific T cell expansion also presents difficulties, as the antigen should be presented in the context of MHC I or II depending on the type of T cell^7^. In part, these issues can be solved by using feeder cells expressing specific antigen or designed antigen-presenting cells (APCs)^6,8^. In the case of feeder/APC the stimulation is more physiological because cells provide additional signals that facilitate stimulation, proliferation, survival and function of the CAR-T cells. But these advantages are negated by the need to use living cells with their inherent safety considerations, and add the undesired extra steps of feeder cell removal^9^.

We reasoned that it would be possible to use the relatively simple structural requirement for functional CAR interaction with its cognate antigen (a mimic of immunoglobulin-antigen interaction) to develop a more manageable methodology for antigen-specific expansion of CAR-T cells^10^. We felt that a promising strategy to obtain high numbers of functional antigen-specific CAR-T cells would be to stimulate CAR-T cells using antigen exposed on cell membrane fragments organized on relatively uniform microcytoshperes that lack a nucleus. It is known that the blebbing of cell membranes caused by cytochalasin B treatment results in the generation of microvesicles surrounded by plasma membrane^11–13^. Importantly, it has been shown that surface molecules present on the original plasma membrane are also displayed on the membrane surface of these vesicles, and in their native membrane-anchored form, with both proper glycosylation and folding^14,15^. We constructed antigen-specific vesicles (AVs) displaying recombinant antigens for a series of CARs, and showed that incubation of these AVs with their cognate CAR-T cells resulted in antigen-specific signaling, stimulation, and proliferation of the CAR-T cell population. In addition, we showed that stimulation with AVs induced CAR-T cell functional maturation and cytotoxic efficacy *in vitro*. Our data suggest that this new antigen-specific protocol for CAR-T cell stimulation and expansion *in vitro* is more efficient than existing approaches, and may facilitate the further development and application of CAR-T therapy.

## Results

### Generation and characterization of fluorescent AVs

For constructing AVs expressing CAR antigens, first HeLa cells (the most common tumor cell line) were employed (**Figure 1A**). Membrane blebbing was induced by treatment of cells with cytochalasin B. Cytochalasin B rapidly disrupts the actin cytoskeleton, resulting in extrusion of the nucleus, detachment of the plasma membrane, and formation of long tubular extensions from the cell^12,16^. These extensions can be sheared off by shaking to form vesicles which carry a replica of surface proteins expressed by the source cell^11^ (**Figure 1 B and C**). We envisioned that AVs could serve as a mimic of tumor cells, since they would possess the same set of surface molecules. Such AVs can bind, and hence, label and identify CAR-T cells of a desired specificity. Moreover, generation of stable tumor cell lines simultaneously expressing a fluorescent protein and a specific CAR target antigen on their surface could provide an unlimited source of CAR-specific AVs suitable for CAR-T cell binding, and therefore, stimulation and detection. In addition, these AV’s could be further stabilized for storage and future use in CAR-T research or therapy.

**Figure 1.**
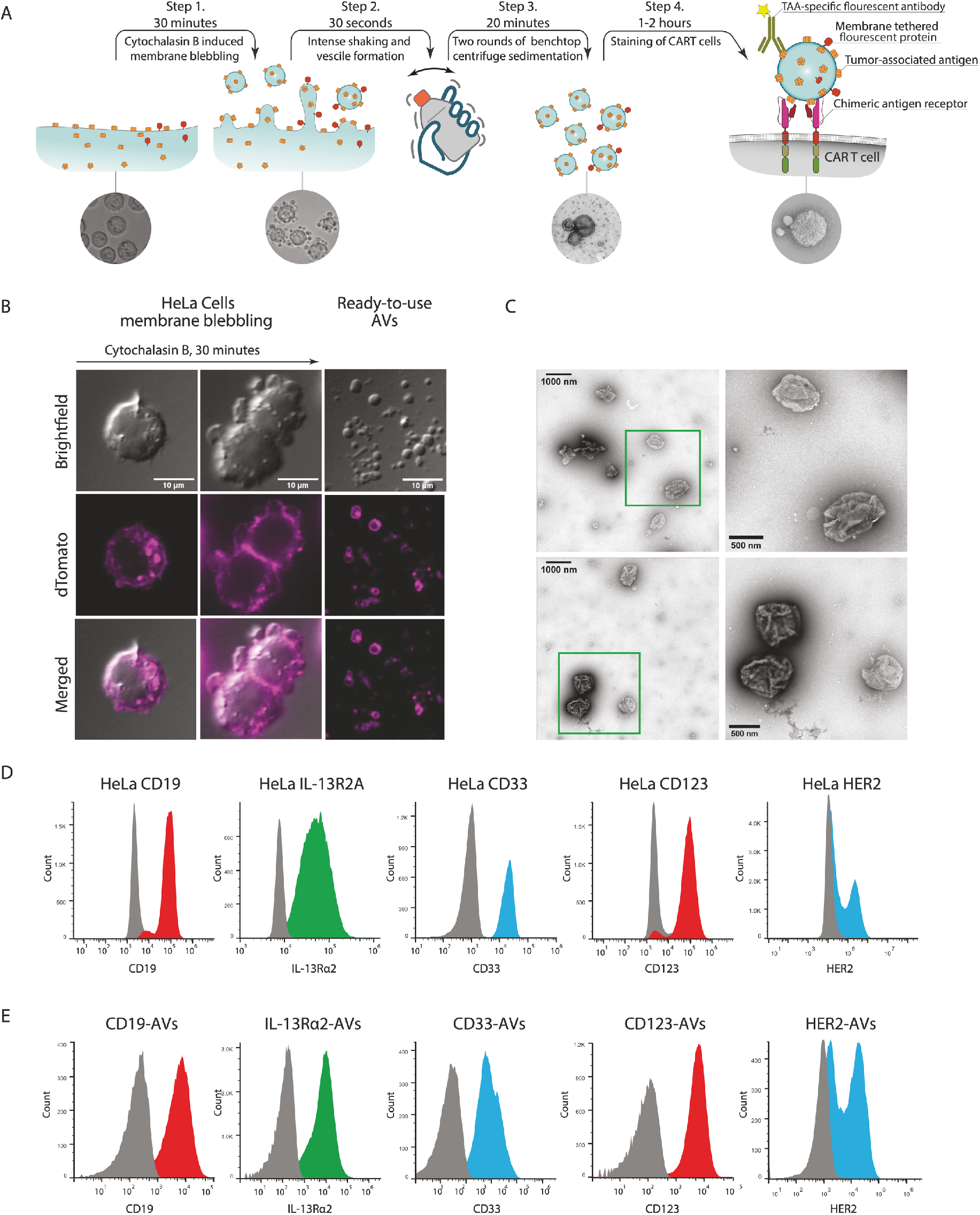
Generation and characterization of fluorescent AVs with surface expression of CAR antigens and dTomato. **(A)** The scheme shows major steps in the AV generation protocol. **(B)** Confocal images of HeLa/dTomato cell membrane blebbing during Cytochalasin B treatment and artificial vesicles after purification. **(C)** TEM images of purified AVs at different magnifications. **(D)** Staining of HeLa/dTomato cell lines with surface expression of CD19, IL-13Ra2, HER2, CD33 and CD123 antigens. Control cells and cells transduced with corresponding lentiviruses were stained with fluorescent antibodies. Stained cells were analyzed by flow cytometry, and results plotted as histograms. Histograms of control (non-transduced) cells are shown in grey. **(E)** Expression of antigens on the surface of AVs. Vesicles were stained with fluorescent antibodies, and analyzed as in (D). Control vesicles obtained from HeLa/dTomato cells which do not have target antigens on cell membrane are shown in grey.

To obtain fluorescent AVs with surface expression of an individual CAR tumor antigen, HeLa cells were transduced stepwise with lentiviruses encoding the membrane-tethered fluorescent protein dTomato^17^ and one of a series of tumor-associated antigens (**Figure 1B and D**). The resulting Hela cells expressing the fluorescent dTomato protein and CD19, CD33, CD123, IL13Ra2, or HER2 (**Figure 1D**) were incubated with Cytochalasin B, then broken apart by shaking. Next, simple centrifugation allowed isolation of ready-to-use suspensions of the antigenic vesicles (AVs) expressing both dTomato protein and one of the CAR antigens – CD19-AVs, IL-13Ra2-AVs, HER2-AVs, CD33-AVs or CD123-AVs, respectively. Flow cytometry of AV suspensions led to an estimation that from each HeLa cell it is possible to obtain approximately 4 AVs. (**Supplementary Fig. 1B and 1C**). The presence of target antigen on the AV surface was confirmed by flow cytometry following immunostaining of AV suspensions with a mAb specific for the corresponding antigen (**Figure 1E**).

To investigate the structure of the AVs, and to confirm that suspensions indeed contained microvesicles), CD19-AVs were analyzed using transmission electron microscopy (TEM, **Figure 1C, Supplementary Fig. 1A**). TEM images demonstrate that the diameter of the AVs ranges from 200 to 2 000 nm.The AVs looked creasy, which is typical for the relatively large artificial vesicles^11^.

### AVs bind specifically to the surface of their cognate CAR-T cells

The presence of antigen on the surface of AVs does not guarantee that they are capable of binding CAR-T cells. Therefore, the ability of AVs to bind specifically to their cognate CAR-T cells (CD19-CAR, IL-13-CAR, HER2-CAR, CD33-CAR and CD123-CAR T cells) was examined. The structures of corresponding CARs and differences in structure/binding with respective antigens are shown in **Figure 2A**. CARs of diverse design, including variations in hinge regions (CD8a-hinge or CH2-CH3 of IgG Fc domain) or in the target-recognition domain (scFvs or IL-13 cytokine) were used in binding/detection experiments (**Figure 2A**). CAR-T cells were stained in a single step by the AVs or by commercially available CAR detection reagents (mAb to hinge regions, CD19-Fc recombinant protein^18^) for side-by-side comparison (**Figure 2B, Supplementary Fig. 2, Supplementary Fig. 3B and 3C, Supplementary Fig. 4A**). The results of these analyses demonstrate that AVs can bind and stain CAR -T cells similarly to mAbs or recombinant proteins. Brightness of CAR-T cells stained with dTomato-labeled AVs and strength of vesicle binding to cell membranes allow for FACS sorting and purification of CAR-T cells from mixtures including CAR-negative T cells. CAR-T cells stained with AVs were successfully sorted from mixtures with other T cells (**Figure 2C**). Sometimes, CAR-detection antibody or recombinant proteins are not available from vendors. In this case, AVs with designed CAR specificity will be very useful for detection of CAR-transduced cells (**Figure 2B and C, Supplementary Fig. 4B**) as it is shown for CD33-CAR and two different variants of CD123-CAR (**Supplementary Fig. 3A**). Thus, AVs do not have limitations associated with variation in CAR structure or in the structure of a target antigen since they bind directly to the exposed antigen-binding region of the CAR.

**Figure 2.**
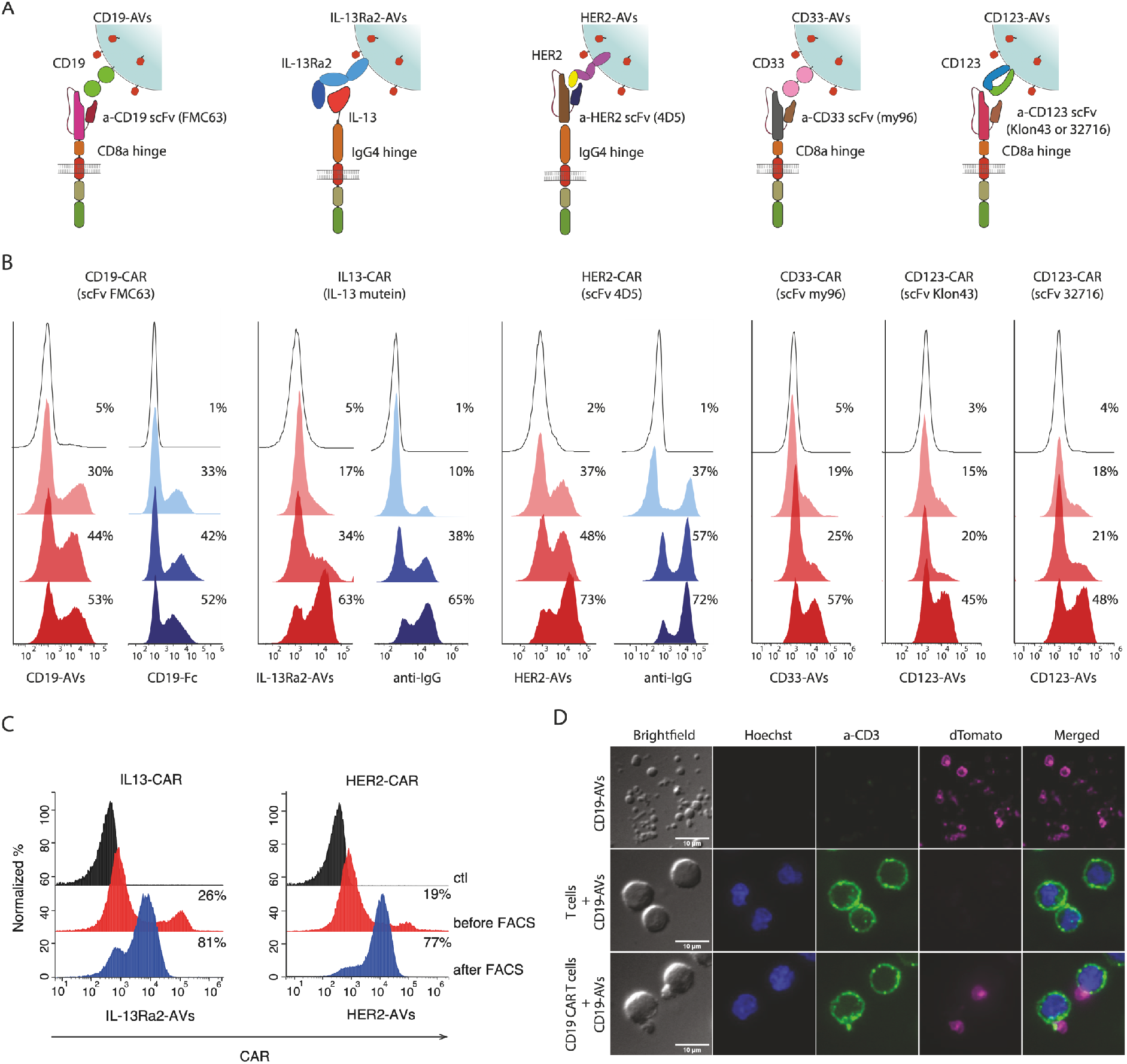
AVs expressing CAR antigens bind CAR-T cells in a specific and dose-dependent manner, and can be used as CAR detection reagents. **(A)** Schematic drawings of interactions between CARs and their target antigens exposed on the surface of AVs along with dTomato. **(B)** Side-by-side comparison of CAR-T detection/staining by the AVs and IgG-specific antibodies or CD19-Fc (CAR detection reagents). Suspensions of cells with different proportions of CAR-T cells (CAR-T cells were spiked with control CAR-negative T cells) were stained in parallel with fluorescent mAbs against CARs or corresponding AVs. Stained cells were analyzed by flow cytometry and plotted as histograms. AVs on the surface of cells were visualized using dTomato fluorescence. Empty black histograms show background signal of CAR-negative T cells. Numbers indicate proportions of detected CAR-T cells. **(C)** IL13-CAR and Her2-CAR detection before and after FACS sorting of CAR-positive population using staining with dTomato-AVs. FACS sorted cells stained with AVs were re-analyzed by flow cytometry using dTomato fluorescence. Black histogram represents staining of control cells. **(D)** Confocal imaging of CD19 AVs, control T cells, and CD19 CAR-T cells. Control T cells or CD19 CAR-T cells were incubated with a suspension of CD19 AVs prior to microscopy. Nuclei were stained with Hoechst 33342 (blue), T cells APC-labeled anti-human CD3 mAb (green). Unbound vesicles and vesicles attached to the surface of cells after incubation were visualized using dTomato fluorescence (magenta). Representative images are shown.

Flow cytometry does not reveal the structure(s) of AVs attached to cells. Are they intact or fragmented? Is the attachment specific or non-specific? To discriminate between these possibilities, confocal microscopy of control cells, AVs and cells following incubation with dTomato-labeled AVs was performed (**Figure 2D, Supplementary Video 1**). One can clearly see that only AVs specific to the CAR, but not control vesicles stay attached to the cell surface. This result confirms that in most cases the binding and fluorescence of CAR-T cells treated with AVs is due to specific attachment of intact AVs to the cell surface.

For additional insight into the interaction of CAR-T cells and AVs, scanning electron microscopy experiments were performed. SEM imaging of control T cells and CD19-CAR cells incubated in parallel with CD19 AVs demonstrated that CD19 AVs do not attach to control T cells, while at least one AV is bound to CD19-CAR T cell (**Figure 3A**). Images show that round-shaped vesicles are glued to the cell surface or sit on top of membrane protrusions. In addition, confocal imaging of CD19 AVs bound to the surface of CD19-CAR-T cells was performed using mAb staining of CD19 antigen on the surface of AVs (**Figure 3B, Supplementary Fig. 3D**). The results of this experiment further confirm that only CD19 AVs binding with CD19-CAR occurs. In the case of control T cells AVs do not attach to the surface of CAR-T cells. Thus, our data on both the microscopic and macroscopic (immunostaining + FACS) level suggest that AVs and cognate CAR-T cell interactions are specific and driven by CAR-antigen interaction.

**Figure 3.**
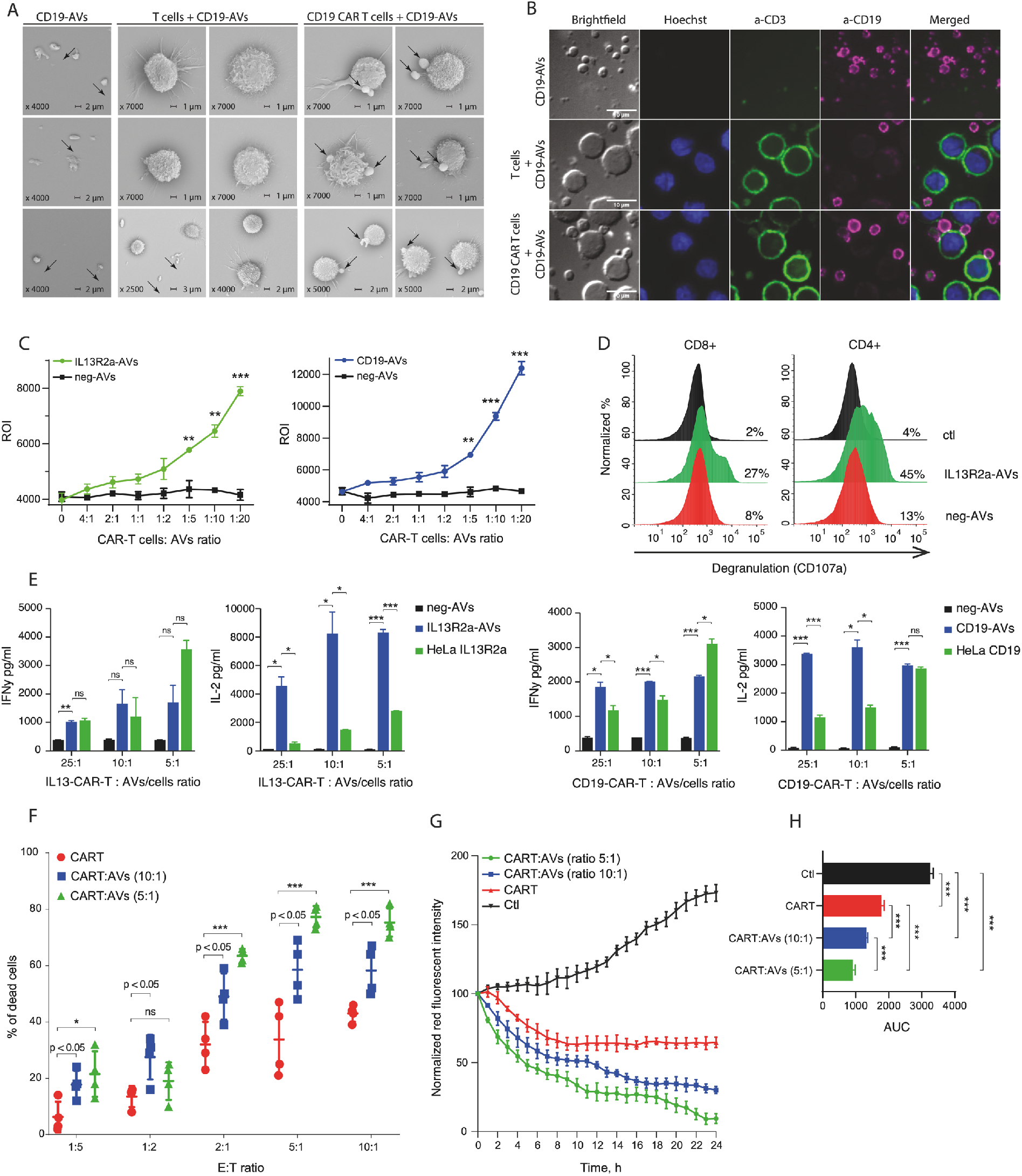
Incubation with AVs stimulates CAR-T cells and enhances their killing activity (cytotoxicity) **(A)** SEM images of the CD19-AVs, control T cells, and CD19 CAR-T cells following incubation with CD19 AVs. Arrows point to AVs attached to the surface of CAR-T cells. **(B)** Mock T cells and CD19 CAR-T cells were incubated with CD19-AVs and stained with anti-CD19-PE antibody. Signal from CD19 mAb is shown in magenta, nuclei were stained with Hoechst 33342 (blue), and cell surface with anti-human CD3 mAb (green). Representative images are shown. **(C)** Analysis of Jurkat-NFAT-luc CAR-T activation after culture with AVs or control antigen-negative vesicles at different CAR-T cell:AV ratios. **(D)** Assesment of IL13 CAR-T cell degranulation in untreated control cells and in cells following culture for 4 hours with IL13R2a-AVs or control antigen-negative vesicles. **(E)** Quantification of IFNγ and IL-2 secretion by IL13-CAR-T or CD19-CAR-T cells co-cultured at various CAR-T:AV ratios with either IL13R2a-AVs or CD19-AVs, or with antigen-negative vesicles. In parallel CD19 CAR- and IL13 CAR-T cells were incubated at various CAR-T:target cell ratios with antigen-expressing cells Hela IL13R2a or Hela CD19, correspondingly. **(F)** *In vitro* killing activity of CD19-CAR-T cells either untouched or cultured with CD19-AVs at 10:1 or 5:1 CAR-T:AV ratio. At day 4 after start of treatment with vesicles, CAR-T cells were incubated with Jeko-1 target cells at various effector-to-target cell ratios. Data are represented as the mean ± s.d. of four experimental replicates and are representative of at least two independent experiments. **(G)** Incucyte killing assay of Jeko-1 by untreated or CD19 AV-treated CD19 CAR-T cells at effector-to-target cell ratio of 3:1. **(H)** Analysis of killing efficacy of CAR-T cells in Incucyte-based killing assay evaluated as Area under curve (AUC) parameter. Statistical analysis was performed using one-way ANOVA with Turkey’s multiple comparisons test. For all panels, ns - not significant, *P < 0.033, **P < 0.002, ***P < 0.001. **(C**,**E**,**F)** Data are presented as the mean ± s.d. of three experimental replicates and are representative of at least two independent experiments. Statistical analysis was performed using unpaired t-test. For all panels, ns - not significant, \**P* < 0.033, \*\**P* < 0.002, \*\*\**P* < 0.001.

### Binding of AVs to CAR-T cells results in activation and proliferation of CAR-T cells

CAR-T cell interaction with target antigen on the cell surface results in stimulation that activates the CAR-T cell. Activation leads to increased proliferation accompanied by production of pro-inflammatory cytokines (IFN-g and IL-2), and secretion of lytic granules containing granzyme and perforin. This sequence of events is a prerequisite for cytotoxicity, the epitome of CAR-T effector function. Therefore, to state that AVs are capable of inducing CAR-T cell function it is necessary to test experimentally if incubation with AVs will cause stimulation, division (and hence expansion) and cytotoxic function (measured as degranulation and lytic activity).

First, the ability of AVs to initiate downstream signaling of the CAR was tested. Two model Jurkat T cell lines with stable expression of NFAT-fluc reporter and different CARs were generated. One of them had surface expression of IL-13-CAR, and another one, CD19-CAR (**Supplementary Fig. 5B**). Binding of the CARs with their target antigens in these cell lines should result in downstream signaling, activation of NF-AT transcription factor and induction of NF-AT-dependent transcription of fluc cDNA. Production of fluc and its enzymatic activity increases upon CAR triggering and directly correlates with the extent of CAR stimulation. Jurkat CAR reporter lines were incubated with corresponding AVs at different ratios, and fluc activity was measured using a luciferase assay. As expected, increasing the number of vesicles per Jurkat-CAR-NFAT-fluc cell led to significant enhancement of normalized luminescence signal (**Figure 3C**). Notably, control vesicles obtained from the parental cell line lacking antigen were not able to induce fluc activity. This result demonstrates the capacity of AVs to trigger CAR-T stimulation.

### Induction of CAR-T functional maturation by the AVs

Next, in order to determine if stimulation of CAR-T cells by AVs induces functional maturation, we analyzed their ability to secrete lytic granules filled with granzyme and perforin. Efficiency of this degranulation process can be assessed indirectly by surface expression of CD107a marker (LAMP-1). CAR-T cells obtained by lentiviral transduction of donor T cells with IL-13-CAR-expressing constructs were incubated with AVs (at 1:1 ratio), then analyzed for the surface level of CD107a on CD4 and CD8 T cells. Our data indicate that AVs expressing IL13Ra2 dramatically increase degranulation of both CD4 CAR-T cells and CD8 CAR-T cells. Control empty microvesicles had minimal effect on degranulation, indicating that this functional maturation was antigen-specific (**Figure 3D**).

To characterize the induction of CAR-T effector function by AVs in more detail, production of pro-inflammatory cytokines, IFN-g and IL-2, was determined using ELISA assay of culture supernatants of IL13-CAR-T or CD19-CAR-T cells following incubation with different ratios of control microvesicles, AVs expressing IL13Ra2/CD19, or HeLa cells expressing IL13Ra2/CD19. Two different combinations of CAR-target antigen were used to minimize potential artifacts (**Figure 3E**). At high ratios of AV:CAR-T AVs were comparable or better than antigen-expressing HeLa cells in terms of cytokine induction in both IL13 and CD19 CAR-T cells. But surprisingly, in both types of CAR-T cells production of IL-2 was much higher at intermediate as well as high AV:CAR-T ratios. Total surface of HeLa cells should be much larger that net surface of equal number of AVs, and surface composition should be also the same or very similar.

Although degranulation and cytokine production correlate with the cytotoxic activity of CAR-T cells, we wanted direct proof of the ability of AV-stimulated CAR-T cells to specifically kill their target cells. We used Jeko-1 cells as the target population and compared cytotoxicity when they we incubated with either untreated or AV-stimulated CD19-CAR transduced T cells (**Supplementary Fig. 5E**). In order to study the effect of AV dose vesicles were added at two different ratios. The results demonstrated clearly that even at a low CAR-T:AV ratio (10:1) the killing activity of CD19-CAR-T is greatly increased by stimulation with AVs. (**Figure 3F**). Incucyte technology was used to further confirm the effect of AVs on killing of target cells. One can see that Incucyte data strongly support the observation that pre-incubation wit AVs potentiates the killing of target cells (**Figures 3G and H**).

### Incubation with AVs results in antigen-specific expansion of functional CAR-T cells

Next, we wanted to determine if AVs could be used for antigen-specific expansion of CAR-T cells. To test this, CD19-positive (CD19-AVs) and control (CD19-negative) AVs were added to CD19-CAR-T cells following CFSE labeling. IL-2 stimulated cells served as a positive control. In this assay the extent of CFSE dilution during the process of cell proliferation is proportional to the number of divisions. CD19-CAR-T cells were stimulated with IL-2 or AVs for 4 days, then CFSE fluorescence was assessed by flow cytometry. As expected, there was no significant expansion of CAR-T cells incubated with control AVs, while the CAR-T cell proliferation rate in the presence of CD19-AVs was increased, and in fact, exceeded the rate observed with IL-2 stimulation (**Figure 4A**). It is likely that strong stimulation of CD4^+^ and CD8^+^ subsets by CD19-AVs led to increased secretion of IL-2 and autocrine IL-2 signaling, accounting for the proliferation rates observed in the IL-2 and CD19-AV treated samples (**Supplementary Fig. 5A**).

**Figure 4.**
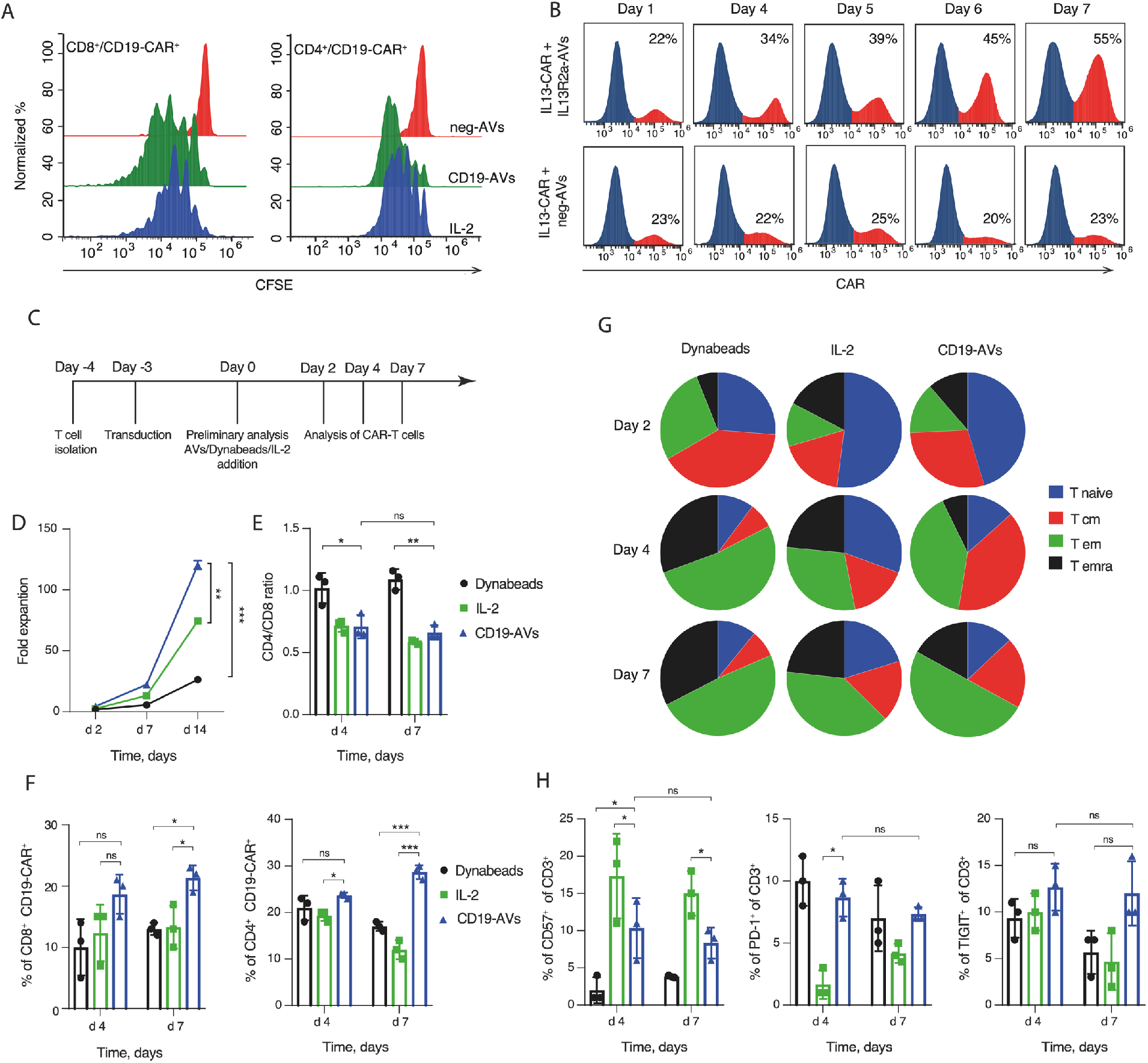
Antigen-specific expansion of CAR-T cells. **(A)** CFSE-based tracking of proliferation in CD19 CAR-T cells that were cultured in the presence of either IL-2, or with antigen-positive (CD19-AVs) or antigen-negative (neg-AVs) artificial vesicles at a 5:1 ratio for 4 days. **(B)** Increase in the proportion of IL13 CAR-T cells in a mixture with CAR-negative T cells that were cultured for 7 days in presence of antigen-positive (IL13R2a-AVs) or antigen-negative vesicles at a 5:1 ratio. Numbers show proportions of CAR-T-positive cells. (**C)** The scheme shows major time points in the expansion experiment protocol. **(D)** Analysis of CD19 CAR-T-cell expansion following exposure to different activation stimuli (Dynabeads, IL-2 or CD19-AVs). **(E)** Assessment of changes in the CD4/CD8 ratio in a subpopulation of CD19 CAR-T cells exposed to different activation stimuli (Dynabeads, IL-2 or CD19-AVs). **(F)** Analysis of changes in CD4^+^ CAR^+^ and CD8^+^ CAR^+^ proportions following exposure to different activation stimuli. **(G)** FACS quantification of changes in the CD19-CAR cell surface phenotype (reflects maturation/differentiation state) in response to different stimulating agents: CD3/CD28 Dynabeads, IL-2 or CD19-AVs. **(H)** FACS quantification of CD19-CAR cells expressing exhaustion markers PD-1, CD57 and TIGIT among CD3^+^ cells in samples cultured with Dynabeads, IL-2 or CD19-AVs. **(D)** Data are presented as the mean ± s.d. of three experimental replicates and are representative of at least two independent experiments. Statistical analysis was performed using unpaired t-test. **(E**,**F**,**H)** Data are presented as the mean ± s.d. of three experimental replicates and are representative of at least two independent experiments. Statistical analysis was performed using one-way ANOVA with multiple comparisons. For all panels, \**P* < 0.05, \*\**P* < 0.01, \*\*\**P* < 0.001, \*\*\*\**P* < 0.0001.

Next, the frequency of T cells expressing IL13-CAR was determined during the course of prolonged incubation with IL13R2a-positive (IL13R2a-AVs) or IL13R2a-negative (control) AVs. Donor T cells were transduced with IL-13-CAR lentiviral particles giving rise to a mixture of CAR-positive and CAR-negative T cells. Following the addition of antigen-expressing or control AVs, the proportion of IL13-CAR-positive and negative cells was measured at days 1, 4, 5, 6 and 7 using immunostaining followed by flow cytometry. In samples stimulated with control AVs the frequency of the CAR-positive subpopulation remained almost constant during the entire 7 day incubation. In contrast, in samples stimulated with IL13R2a-AVs the proportion of CAR-positive cells increased drastically from 22% at day 1 to 55% at day 7 (**Figure 4B**). Taken together, these data demonstrated the effectiveness of antigen-expressing AVs to specifically stimulate and support the division and accumulation of their cognate CAR-T cells in a mixture with unmodified lymphocytes.

To confirm that CAR-T cell populations expanded using our AV methodology do not become terminally differentiated effector cells that often suffer from functional exhaustion, the surface phenotype of freshly isolated T cells transduced with CD19-CAR and stimulated with anti-CD3/CD28 Dynabeads, IL-2, or CD19-AVs on day 3 post-transduction was assessed using immunostaining and FACS analysis. (**Supplementary Fig. 5D, Figure 4C**). CD3/CD28 Dynabeads and IL-2 are routinely used to expand CAR-T cells after viral transduction of donor T cells, and were chosen to perform a direct comparison with stimulation by CD19-AVs. Re-stimulation with Dynabeads or IL-2 to provide alternative stimulation signals) is currently widely used in expansion protocols for stimulation of therapeutic T cells^1^. Panels of fluorescent mAbs were selected to discriminate between different effector and memory or memory-like populations. In addition, the level of T cell exhaustion markers on the membrane of expanded cells was measured using corresponding antibodies and flow cytometry of stained cells. Prior to stimulation (day 0), the CD4^+^/CD8^+^ ratio, expression of lymphoid homing molecules CD62L/CD45RA, and the basal exhaustion level of T cells were determined (**Supplementary Fig. 5C and 5**F). T cells expanded under different conditions were highly viable. We were gratified to learn that at day 7 the expansion rate of the AV-stimulated CAR-T cells was five times higher than the rate observed using the Dynabeads protocol. (**Figure 4D**). While balanced (equal) CD4^+^ and CD8^+^ T cell expansion was observed with Dynabeads stimulation, AV or IL-2 stimulation resulted in a CD8^+^ expansion rate was higher at days 4 and 7 than the expansion rate for CD4^+^ T cells (**Figure 4E**). Moreover, a substantial increase of the CAR-positive population was observed in both the CD4^+^ and CD8^+^ T cell subsets treated with AVs (18% to 21% for CD8^+^ and 23% to 29% for CD4^+^ T cells). This expansion of CAR-positive populations was significantly higher than that seen under other tested conditions (**Figure 4F**). An increase in the CAR-positive population was also observed in similar experiment with autologous CAR-T product (**Supplementary Fig. 6D**). While re-stimulation with Dynabeads led to a prevalence of effector memory (T_em_) and terminally differentiated T cells (T_emra_), incubation with AVs resulted in fewer T_emra_ and a higher frequency of T_cm_ (central memory) and T_em_, cells, even in comparison with IL-2 treatment of the healthy donor CAR-T cells (**Figure 4G**). To further assess the potential clinical usefulness of our AV expansion methodology in CAR-T therapy, the effect of AVs on the generation of autologous CAR-T product ex vivo was tested (**Supplementary Fig. 6A**). When compared with controls, the population of CD19-CAR-T cells expanded using CD19-AVs retained a more naïve state, with a higher frequency of naïve and central memory T cells and enhanced killing activity. (**Supplementary Fig. 6B and 6C**). Despite a nearly fivefold greater expansion boost at day 7 compared to Dynabeads stimulation, AV stimulation resulted in no sizable difference in the frequency of T cells expressing exhaustion markers PD-1 and TIGIT (**Figure 4H**). We also analyzed changes in the proportion of CD57-positive cells throughout the stimulation process. CD57 is a marker of cytotoxicity expressed on NK and activated CAR-T cells, and at baseline positive cells comprised approximately 1% of the population. After supplementation with IL-2 or AVs the proportion of CD57+ T cells was markedly increased - up to a value of 15% or 10% at days 4 and 7 respectively) (**Figure 4H**).

## Discussion

The expansion of antigen-specific populations of T cells is a critical but challenging step in adoptive T cell therapy^19^. This holds true for CAR-T cell-based technologies which are increasingly being used to treat hematological malignancies, but whose application to other forms of cancer has not yet met with wide success. In CAR-T cell therapy relatively small numbers of cytotoxic CAR-transduced donor T cells must be expanded *ex vivo* to obtain numbers adequate to mount an effective *in vivo* response against the target malignancy. Control of outgrowth of CAR-negative T cells is becoming increasingly important in the case of modern, safe, but relatively ineffective non-viral ways of CAR delivery such as piggyback and sleeping beauty transposons^20^ During the expansion process, careful evaluation of the proportion of antigen-specific CAR-T cells, as well as their functional fitness, is needed. Minimizing contamination of the therapeutic T cell population by expansion of non-therapeutic T cells co-existing in the original population, or by replication-competent target cells is increasingly seen as a key to the overall success of CAR-T therapy. Both of these complications are overcome by stimulating with AVs. AVs only stimulate their cognate T cells, thereby avoiding expansion of co-existing non-therapeutic T cells. And AVs as vesicles derivative of the plasma membrane of target cells, have all the advantages of living target cells without the complication of replication. Not surprisingly, the length of *ex vivo* time required in the processing of therapeutic CAR-T cells has also been shown to be a significant factor in treatment success - the less time required for ex vivo T cell expansion, the more likely the population maintains its cytotoxic properties and non-terminal differentiation state^4^. Restricted stimulation time and a large proportion of CAR-T cells in effector-memory differentiation state safeguard the population from functional exhaustion^6^.

Currently, the main approaches to expand and propagate CAR-T cells are 1) incubation with IL-2 and other cytokines that support survival and proliferation of T-cells, and 2) strong non-specific stimulation with anti-CD3/CD28 monoclonal antibodies^6,9^. The later takes advantage of the presence of natural αβ TCR along with recombinant CAR on the membrane of CAR-T cells. This signal differs from the signals provided by the CAR or specific stimulation of natural TCR with its cognate antigen in the context of MHC presentation^7^. To deliver the CD3/CD28 signal, corresponding mAbs are attached to the surface of polymer beads or plastic tissue culture plates. Since rigid immobilization of mAbs on a polymer makes their mobility in the membrane following cross-linking of the TCRs impossible, the signal they provide is not optimal in comparison to antigenic stimulation. On the other hand, antigen-specific stimulation using antigen-presenting cells/feeder cells provides extra co-stimulation signals and secretion of cytokines that favor better quality stimulation in terms of effector function^2^.

To combine the advantages of antigen-specific stimulation and feeder cells, we proposed to generate extracellular vesicles expressing the CAR target antigen on their surface, and use them for CAR-T stimulation, activation, expansion, and finally, *in vitro* staining and purification. The idea was that extracellular vesicles would mimic the target cell membrane microenvironment and provide antigen-specific stimulation by exposing high amounts of target recombinant protein on their surface. We termed these constructs Antigenic Vesicles (AVs), and our protocol for their rapid, simple, robust and inexpensive generation is presented in this article. This protocol utilizes the advantage of cytochalasin B treatment and stabilization of AVs by Pluronic. AVs lack nuclei and cytoskeleton, are stable in solution and can be used to detect CAR-T cells *in vitro*. To simplify the use of AVs for CAR-T cell detection, bright fluorescent protein dTomato was overexpressed along with one of the target antigens in HeLa or HEK293 cell lines. AVs are relatively large in size in comparison to natural extracellular vesicles, and we showed that they express high amounts of target antigen on their surface. AVs were non-toxic, and even at low concentrations were capable of strong stimulation of CAR-T cells (CD19 CAR and IL13 CAR). This stimulation led to the generation of effector CAR-T cells that were highly viable, proliferated, and secreted proinflammatory cytokines (IL-2 and IFN-g), granzyme and perforin. In other words, AV-stimulated CAR-T cells were highly activated and functional according to all indirect tests. To further prove that AV treatment produces cytotoxic CAR-T cells we carried out functional experiments which demonstrated that AV-stimulated cells were capable of killing target Jeko-1 cells.

We felt our work would be incomplete without direct comparison of AV stimulation with the traditional stimulation methods using IL-2 or CD3/CD28 magnetic beads. Surprisingly, AV stimulation of CAR-T cells was at least comparable, but in many cases more potent, than stimulation with either IL-2 or Dynabeads. Our work suggests that AV-stimulation has a number of advantages over stimulation methods currently being used. First, in mixtures of CAR-positive and CAR-negative T cells, AV-stimulated CAR-positive T cells divided faster, so that the proportion of CAR-positive cells increased, in some cases more than doubling after 7 days of stimulation. This proliferative boost accounted for a more than 5-fold increase in the expansion of CAR-positive cells in comparison to CD3/CD28 stimulation, and a 1.5-fold increase in comparison to IL-2 stimulation. Second, effector CAR-T cells resulting from AV stimulation were highly functional, and in multiple tests their capacity to secrete cytokines, degranulate, and kill target cells was shown to be as good or better than that of cells obtained by IL-2 or CD3/CD28 stimulation. Third, the CD4/CD8 ratio is a property that has recently been shown to be important for the function of the adoptively transferred T cells^21^, and AV stimulation resulted in less significant changes in the CD4/CD8 ratio compared with traditional approaches. Fourth, AV stimulation was also superior in terms of generation of higher proportions of central memory and effector-memory populations, as well as a smaller proportion of functionally exhausted cells. Thus, these parameters, in addition to the simple and inexpensive work flow, suggest that the use of AVs for the *ex vivo* activation and expansion of CAR-T cells has the potential to significantly advance adoptive T cell therapy. We believe multiple factors may contribute to the surprising effectiveness of antigen-loaded vesicles in the stimulation and expansion of CAR-T cells. Perhaps it is due to a difference in the quality of signal(s) transduced through the chimeric receptor in comparison to IL-2 and anti-CD3/CD28. Additionally, the size and plasticity of the AV plasma membrane may facilitate T cell activation, and/or there may be other molecules on the surface of AVs that deliver additional CAR stimulation signals. Finally, AVs may be frozen, stored, and standardized for use in multiply patients with similar disease.

Lastly, we identified imaging as another potential use of AVs by showing that fluorescent protein-labeled AVs could be used to detect CAR-T cells. These fluorescent AVs were bright, and in a rapid, dose-dependent, and CAR-antigen specific-manner bound to their cognate CAR-T cells. Moreover, it was possible to sort AV-stained CAR-T cells using FACS sorting. In these results fluorescent AVs were comparable to mAbs or commercially available CAR detection reagents. However, fluorescent AVs did show an advantage in that several CARs that could not be detected using commercial CAR detection reagents, were successfully detected on the surface of CAR-T cells following incubation with fluorescent AVs. Also, it is possible to make AVs of dual specificity that could be used to detect different CARs. In summary, the presented protocol to generate AVs using Cytochalasin B treatment of antigen-expressing cell lines is a simple and universal approach to make a reagent for *ex vivo* stimulation and expansion, as well as detection of CAR-T cells of desired specificity. We speculate that AVs will significantly improve the development of products for adoptive T cell therapy in the near future.

### Experimental Section/Methods

#### Cells and culturing conditions

Cell lines were cultured in media (DMEM or RPMI 1640) supplemented with 10% FBS (Gibco), 10 mM HEPES, 100 U/ml penicillin, 100 ug/ml streptomycin, and 2 mM GlutaMAX (Gibco). The HEK293T lentiviral packaging cell line (Clontech) and HeLa were cultured in DMEM (Gibco). The HeLa and Jeko-1 cell lines were obtained from the Institute of Cytology RAS culture collection (St. Petersburg, Russia). Jeko-1 cell line was transduced with NucLight Red Lentivirus reagent (Sartorius) to generate stable line expressing red fluorescent protein mKate2. Human peripheral blood mononuclear cells (PBMCs) were isolated from the blood of patients and healthy donors by gradient density centrifugation on a Ficoll-Paque (GE Healthcare) according to standard protocol. All patients and healthy donors provided informed consent. The cell line was repeatedly tested for the presence of mycoplasma contamination (MycoReport Mycoplasma Detection Kit, Evrogen, Russia). Jeko-1 and human T-cells were cultured in RPMI (Gibco).

#### Flow cytometry analysis

The following fluorophore-conjugated mAbs were used: anti-human CD33 APC (Biolegend), anti-human CD123 PE (Biolegend), anti-human CD19 PE (Biolegend), anti-human IL13Ra2 FITC (R&D Systems), anti-human HER2 APC (R&D Systems), anti-human CD8-PE-Cy7 (BD Biosciences), anti-human CD4-APC-Cy7, anti-human CD107a-PE (BD Biosciences). The CAR molecules were detected by staining with CD19 CAR Detection Reagent (Biotin) (Miltenyi Biotec), FITC-Labeled Human CD19 (Acro Biosystems), goat anti-human IgG antibody conjugated with DyLight650 (Thermo Fisher Scientific) or recombinant protein L conjugated with FITC (Acro Biosystems), and streptavidin-APC/Cy7 (Abcam). The AVs surface proteins were detected as follows: the AVs were incubated with monoclonal antibodies or isotype controls for 30 min at 4 °C. Subsequently, the AVs were washed once with 500 µL PBS and analyzed by flow cytometry. NovoCyte 2060 (ACEA Biosciences) instrument was used for flow cytometry, data were analyzed with FlowJo X10 (FlowJo) and NovoExpress Software (ACEA Biosciences). Cell lines were sorted by SH800 Cell Sorter (Sony). *Chimeric antigen receptors*: The nucleotide sequence of the CD19-CAR^22^ consists of the following parts: anti-CD19 scFv (FMC63), CD8-α hinge region, 4-1BB co-stimulatory domains, and CD3ζ (signaling domain). Synthetic CAR19 gene was purchased (GeneCust) and cloned into the pLV2 lentiviral vector (Clontech) under control of the EF1a promoter. The nucleotide sequences of the previously published CD33 scFv (clone my96^23^) and CD123 (clone Klon43 or 32716^24^) scFv were synthesized and inserted into the pLV2 CAR backbone to replace anti-CD19 scFv. To obtain IL-13-CAR construct, the DNA fragment coding for the IL-2 leader peptide, IL13 mutein (E13Y), human IgG4-CH2CH3 domain, CD28 transmembrane domain, and costimulatory domains of CD28, OX-40, and the CD3ζ-chain was synthesized (GeneCust) as a single cDNA and cloned into the pLV2 lentiviral vector (Clontech) as described for CD19 CAR. The nucleotide sequence of the previously published HER2 scFv (clone 4D5^25^) was synthesized and cloned into the IL-13-CAR backbone.

#### Lentiviral particles and transduction of T cells

The lentiviral particles containing CD19-CAR, IL-13-CAR, CD33-CAR and CD123-CAR were prepared by PEI co-transfection of HEK293T cells with the corresponding lentiviral CAR plasmids and the packaging plasmids. Supernatants containing virus were collected at 48 h post transfection. The titer of lentiviral particles was determined using Lenti-X p24 ELISAs (Clontech). Peripheral blood of patients and healthy donors was obtained from Dmitry Rogachev National Research Center of Pediatric Hematology, Oncology and Immunology. Dynabeads Untouched Human T Cells Isolation Kit (Invitrogen) was utilized for isolation of T cells from human PBMCs. Human T cells were activated with CD3/CD28 Dynabeads (Thermo Scientific) at a 1:1 ratio (Life Technologies) in a complete RPMI containing 40 IU/ml recombinant IL-2 (Pan Biotech) for 24 hours. Activated T cells were re-suspended at concentration of 0.5×10^6^/ml in lentiviral supernatants of CD19-CAR, IL13Ra2-CAR, CD33-CAR or CD123-CAR, then 1 ml of fresh RPMI media supplemented with 40 IU/ml IL-2 was added and cells were plated to 6-well plates. Plates were centrifuged at 1200x g for 90 minutes at 32°C and incubated for 6 hours at 37°C. Then culture medium was changed every 2 days and cells were grown in flasks at a density of 1.0×10^6^/ml.

#### Expression of membrane antigens in HeLa cells

HeLa cells with stable, simultaneous expression of the membrane-tethered fluorescent protein dTomato^16^ and a target antigen (CD19, IL13Ra2, CD33, CD123 or HER2) were obtained by stepwise lentiviral transduction using the lentiviral vector pLV2 (Clontech, CDSs synthesized by GeneCust).

#### Generation of antigenic vesicles (AVs)

AVs were prepared from HeLa cells expressing both the dTomato protein and a CAR antigen using an optimized Cytochalasin B vesicle formation protocol^12^. Cells were trypsinized, washed, resuspended in 10 ml of Vesiculation Buffer (VB: DMEM, 10% FBS, 1% Pluronic F-127) at 1×10^6^ cells/ml and transferred to a T25 flask (Corning, USA). Treatment with Cytochalasin B (Sigma, USA) 10 µg/ml for 30 min at 37°C, 5% CO2 was followed by intensive shaking for 30 sec. The resulting suspension was centrifuged twice (4°C, 50g for 5 min). After the pellets were discarded, the AVs were collected from the supernatant by centrifugation (4°C, 5000g, 20 min), resuspended in VB, and quantified by amount of total protein (BCA reagent, Thermo).

#### CAR-T cell detection using AVs

CAR-T or donor T cells were resuspended in VB at 0.5×106/ml. Cells were incubated with 20 µg (∼ 5 AVs to 1 T-cell) of freshly isolated AVs for 30 min at 4°C, washed three times with 500 µl VB, and then analyzed by flow cytometry.

#### SEM Imaging

After exposure to the AVs in serum-free media, cells were brought into Petri dishes with coverslips (SPL Lifesciences, South Korea) at concentration 50000 cells/slip and incubated for 30 minutes at 37°C, 5% CO2. Following the fixation with 2.5% GA in PBS for 2 hours at room temperature, gradual dehydration in 10%, 30%, 50%, 70%, 80%, 90% and 96% ethanol was performed. Dehydrated samples were twice incubated in 200 µl of HMDS (REACHEM, Russia) for 5 minutes at room temperature and once - in 100 µl of HMDS until the liquid evaporated. The samples were sputter-coated with a 20nm gold layer using Eiko IB 3 ion coater (Japan) and examined with TM3000 microscope (Hitachi, Japan) at 15kV. The same protocol was used to visualize vesicles without cells.

#### TEM Imaging

The carbon-coated TEM grids (Ted Pella, USA) were treated using a glow discharge device Emitech K100X (Quorum Technologies, Great Britain) to hydrophilize the carbon surface and increase the adsorption. The vesicles were deposited onto the grids for 3 min, contrasted with 1% uranyl acetate and dried. Imaging was carried out using a JEM-1400 (Jeol, Japan) transmission electron microscope at 100 kV.

#### Confocal microscopy

The AVs, T cells or CAR-T cells were stained with anti-CD3 antibodies, Hoechst 33342, and anti-CD19 antibodies and applied to poly-L-lysine-covered coverslips, centrifuged for 10 minutes, 100 g at RT. The attached cells were fixed in 4% formalin for 1 hour at room temperature and stained with anti-human CD3 APC/Cy7 (Biolegend), Hoechst 33342 (Sigma), and anti-human CD19 PE antibodies. Confocal images were captured using an Axio Observer Z1 microscope (Carl Zeiss, Jena, Germany) with a Yokogawa spinning disc confocal device (CSU-X1, Yokogawa Corporation of America, Sugar Land, TX, USA).

#### CAR-T cell activation assay

Reporter cell line Jurkat-NFAT-Lucia (Invivogen, France) expressing CD19-CAR (Jurkat CD19-CAR) or IL13-CAR (Jurkat IL13-CAR) were obtained by lentiviral transduction as described before. For activation assay 5×10^4^ Jurkat CAR-T cells were mixed with freshly isolated antigen-positive or antigen-negative AVs at CAR-T cell-to-AVs cell ratios (0, 4:1, 2:1, 1:1, 1:2, 1:5, 1:10 or 1:20) in a 96-well plate and incubated for 24 hours at 37°C. 25µl of supernatant was separated from cells by centrifugation (4°C, 300g, 5 min) and transferred to the opaque 96-well plate. Activation of reporter Jurkat-NFAT-Lucia CAR-T cells was measured by level of luciferase activity following reaction with Quanti-luc Gold substrate (Invivogen, France) according to the manufacturer’s instructions.

#### T cell degranulation assay

IL13-CAR T cells were resuspended in TexMACS media in concentration 2×10^6^ cells/ml. 100 µl of cell suspension was transferred to the 96-well plate and 2µl of anti-human CD107a antibody (BD) were added in each well. CAR-T cells were mixed with either antigen-positive or antigen-negative AVs at ratio 1:1, or kept untreated (ctl) and were incubated for 2 hours on 4°C. Cell suspension was washed twice with TexMACS media (4°C, 300g, 5 min), stained with anti-human CD3/CD4/CD8 antibodies and analyzed by flow cytometry.

#### CAR-T Cell Cytokine Detection

For cytokine release assays 5×10^4^ CD19-CAR or IL13Ra2-CAR cells/well were incubated with freshly isolated antigen-positive or antigen-negative AVs at CAR-T cell-to-AVs ratios 25:1, 10:1 or 5:1 for 24 h in a 96-well plate in absence of IL-2. As positive control, CD19-CAR and IL13Ra2-CAR T cells were mixed with HeLa CD19 and HeLa IL13R2a cells respectively in CAR-T cell-to-target cells ratios 25:1, 10:1 or 5:1 for 24 h in a 96-well plate in the absence of IL-2. Basal levels of IFN-g and IL-2 were detected in CAR-T samples which were not subjected to stimulation. The supernatant was separated from cells by centrifugation (4°C, 300g, 5 min), transferred to the new plate and stored at -20°C prior use. IL-2 and IFN-γ secretion by human CAR-T cells were analyzed with cytokine-specific ELISA kits (Vector-best, Russia) according to the manufacturer’s instructions.

#### Cytotoxity assays

Prior to cytotoxity assay 5×10^6^ CD19-CAR T cells were incubated with CD19-AVs at T cells-to-AVs ratios of 10:1 or 5:1 for 4 days. To assess the basal level of CAR-T cells cytotoxity, CD19-CAR-T cells were incubated in the absence of AVs. On the day 4 CAR-T cells were collected by centrifugation (4°C, 300g, 5 min) and washed twice with PBS prior to counting.

For LDH-release assay 5×10^3^ Jeko-1 red (NucLight Red) cells per well were mixed with AVs-treated and control CAR-T cells at CAR-T-to-target cell ratios of 10:1, 5:1, 2:1, 1:2 and 1:5 and cultured for 12h in complete RPMI 1640 media supplemented with human IL-2 (30 U/ml). The cytotoxicity of CAR-T cells was evaluated in a standard LDH release assay (CytoTox 96 Non-Radioactive Cytotoxicity Assay, Promega) following the manufacturer’s recommendations.

For Incucyte killing assay 3×10^4^ Jeko-1 red (NucLight Red) target cells were combined with 9 x 10^4^ of AVs-treated and control effector cells per well of 96-well plate and cultured for 24 hours at 37°C. As negative control, 3×10^4^ target cells were plated without addition of effector cells. Changes in fluorescence were monitored every hour with IncuCyte Live-Cell Analysis System (Essen BioScience). Normalized target cell numbers were generated by dividing target numbers after killing to cell counts at the beginning (0 hours) of experiment. To calculate the cytolytic activity of CAR T cells against target cells over time, the area under the curve (AUC) was calculated.

#### T Cell Proliferation Assay

On the day 4 after lentiviral transduction 2×10^6^ IL13-CAR T cells were rested for 48h and then labeled with 5 mμ CellTrace CFSE reagent (Invitrogen) for 5 min on 37°C. Cells were washed twice with PBS and 5×10^5^ CFSE-labeled CAR-T cells were mixed with freshly isolated antigen-positive or control AVs at 1:1 ratio. As a positive control, CFSE-labeled CAR-T cells were cultured in presence of 30 u/ml IL-2. AVs-treated cells were cultured for up to 1 week, culture medium was replaced with fresh medium containing AVs (same amount as at the start of incubation) every 48 h. At the day 4 of co-incubation cells were stained with anti-human IL13 FITC (R&D Systems), anti-human CD4 APC (Biolegend) and anti-human CD8-PE (Biolegend) antibodies and analyzed for proliferation by flow cytometry.

#### In vitro T-cell expansion

At day 4 after lentiviral transduction 1×10^6^ CD19-CAR T cells were stimulated with CD3/CD28 Dynabeads (Thermo Scientific), or 30 U/ml IL-2, or 2×10^5^ CD19-AVs, and cultured for up to 7 days (37C, 5% CO_2_). Fresh culture medium was added following cell counting to keep the cells at a density not exceeding 2.5×10^6^ /ml. Control unstimulated samples were cultured in complete RPMI without IL-2. CD3/CD28 Dynabeads (ThermoFisher Scientific) were used according to the manufacturer protocol. At days 2, 4 and 7 cells were counted, fold expansion was calculated by dividing cell counts at different time points to cell counts at the beginning of experiment. At days 4 and 7 cells were stained with CD19-Fc FITC (R&D Systems), CD4 APC (Biolegend) and CD8-PE (Biolegend), CD45RA-PE (Biolegend), CD62L-APC (Biolegend), CD57-PE-Cy7(Milteni), TIGIT-PE-Cy7 (Milteni), and PD-1-APC (Milteni) antibodies (all anti-human), and analyzed by flow cytometry to assess differentiation status and exhaustion markers of CAR-T cells.

#### Statistics

Statistical analysis was performed using GraphPad Prism 8.0 (GraphPad). Each figure legend denotes the statistical test used. Mean values are plotted as bar graphs, error bars indicate s.d. unless otherwise stated. ANOVA multiple-comparison *p*-values were calculated using Tukey’s multiple-comparisons test. All t-tests were two-sided unless otherwise indicated. For all figures, * indicates P < 0.033, ** indicates P < 0.002, *** indicates P < 0.001.

## Authorship Contributions

U.V.M., R.Y.P., P.D.S., P.N.A., Y.I.A., S.A.I., B.D.V., S.V.O., M.O.V. performed experiments, analyzed and interpreted data, and wrote the paper; T.S.S., K.R.S., K.E.A., M.E.G., O.A.L, Z.M.A., D.S.M., X.J. analyzed and interpreted data and revised the manuscript; G.A.G., M.M.A., S.A.V. and R.A.L designed research, analyzed and interpreted data, and wrote the paper.

## Acknowledgements

Construction of chimeric antigen receptors, lentiviral vector production and transduction of T cells was supported by Russian Scientific Foundation project No. 17-74-30019; Expression of membrane antigens in HeLa cells and construction of artificial vesicles supported by the Russian Foundation for Basic Research grant No. 19-29-04087_mk; The TEM measurements were performed at the User Facilities Center “Electron microscopy in life sciences” at Lomonosov Moscow State University. SEM measurements were carried out using the equipment purchased on account of the Lomonosov MSU Development Program.

---

**Supplementary Figure 1.**
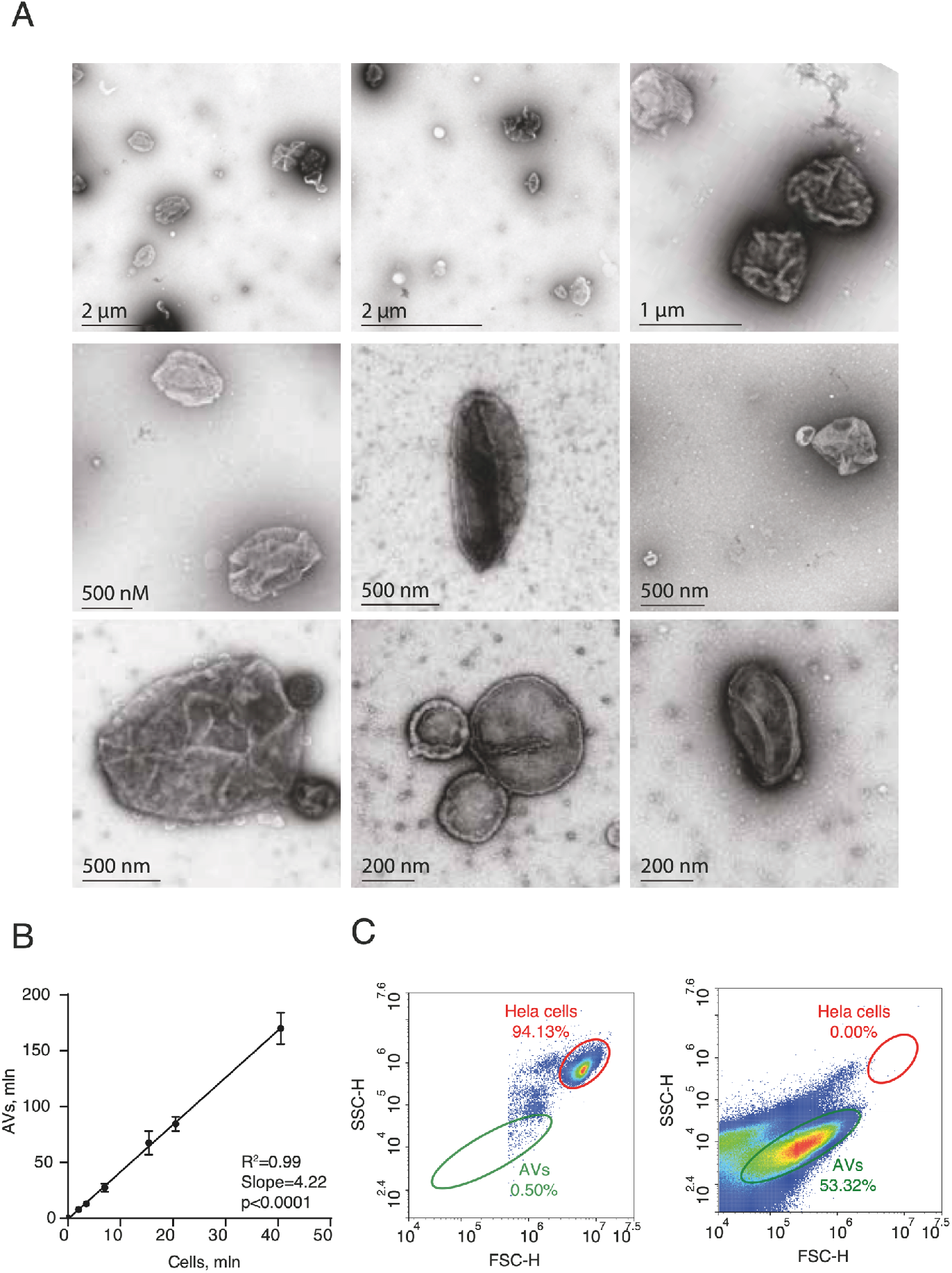
Characterization of isolated AVs. (A) The TEM images of purified AVs (B) Quantification of the isolated AVs. The graph represents the ratio of the number of Hela cells and vesicles isolated from them. Statistical analysis was performed using linear regression model. (C) AVs were separated from Hela cells and analyzed by flow cytometry and plotted as two-dimensional dot plots.

**Supplementary Figure 2.**
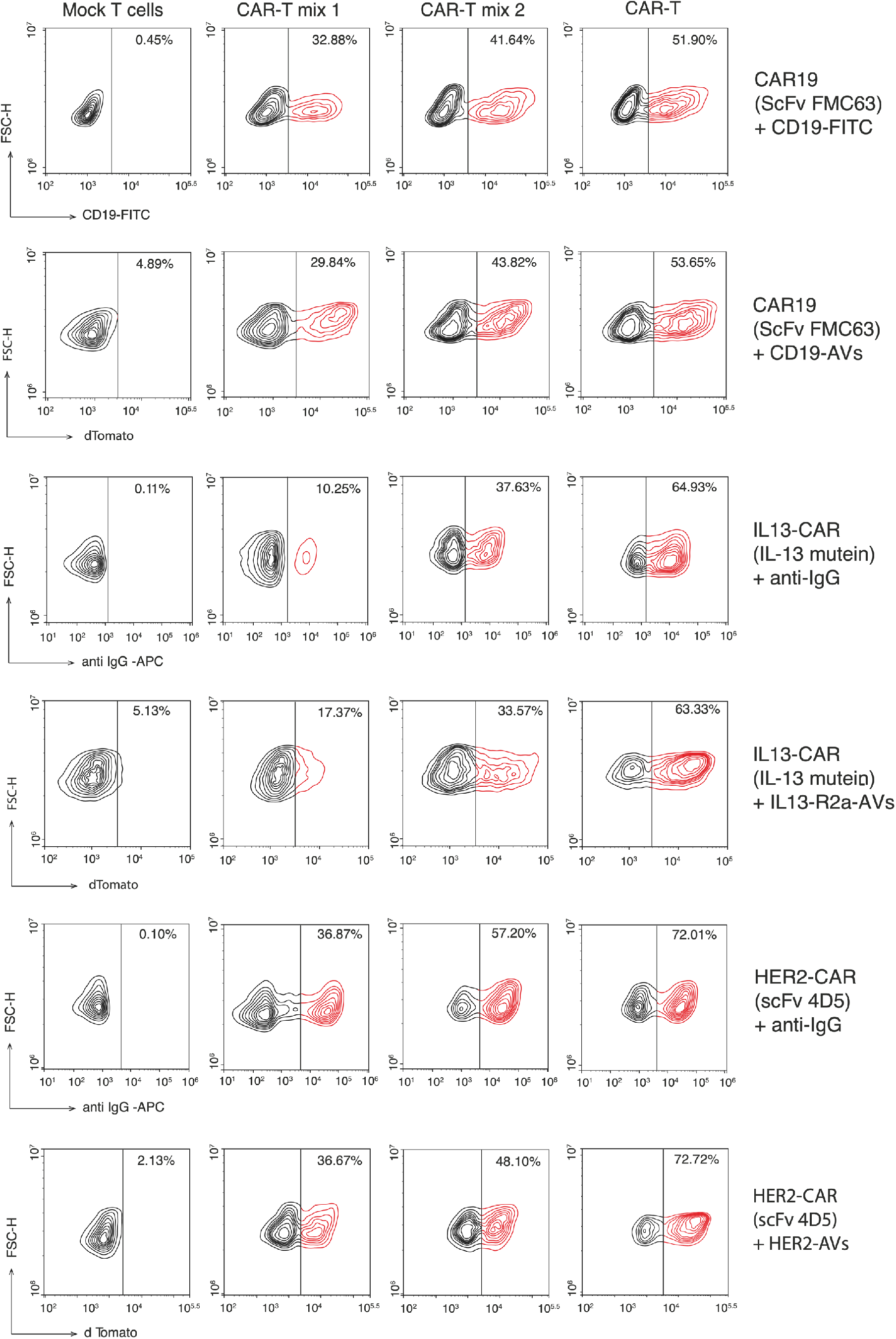
CD19, IL13, and HER2 CAR-T cell detection by the AVs. CAR-T detection by the AVs in comparison to the IgG-specific antibodies or CD19-Fc plotted as two-dimensional contour plots. CAR-T mix 1 and CAR-T mix 2 demonstrate suspensions of cells with different proportions of CAR-T cells (CAR-T cells were spiked with control T cells). Numbers indicate the percentage of detected CAR-T cells.

**Supplementary Figure 3.**
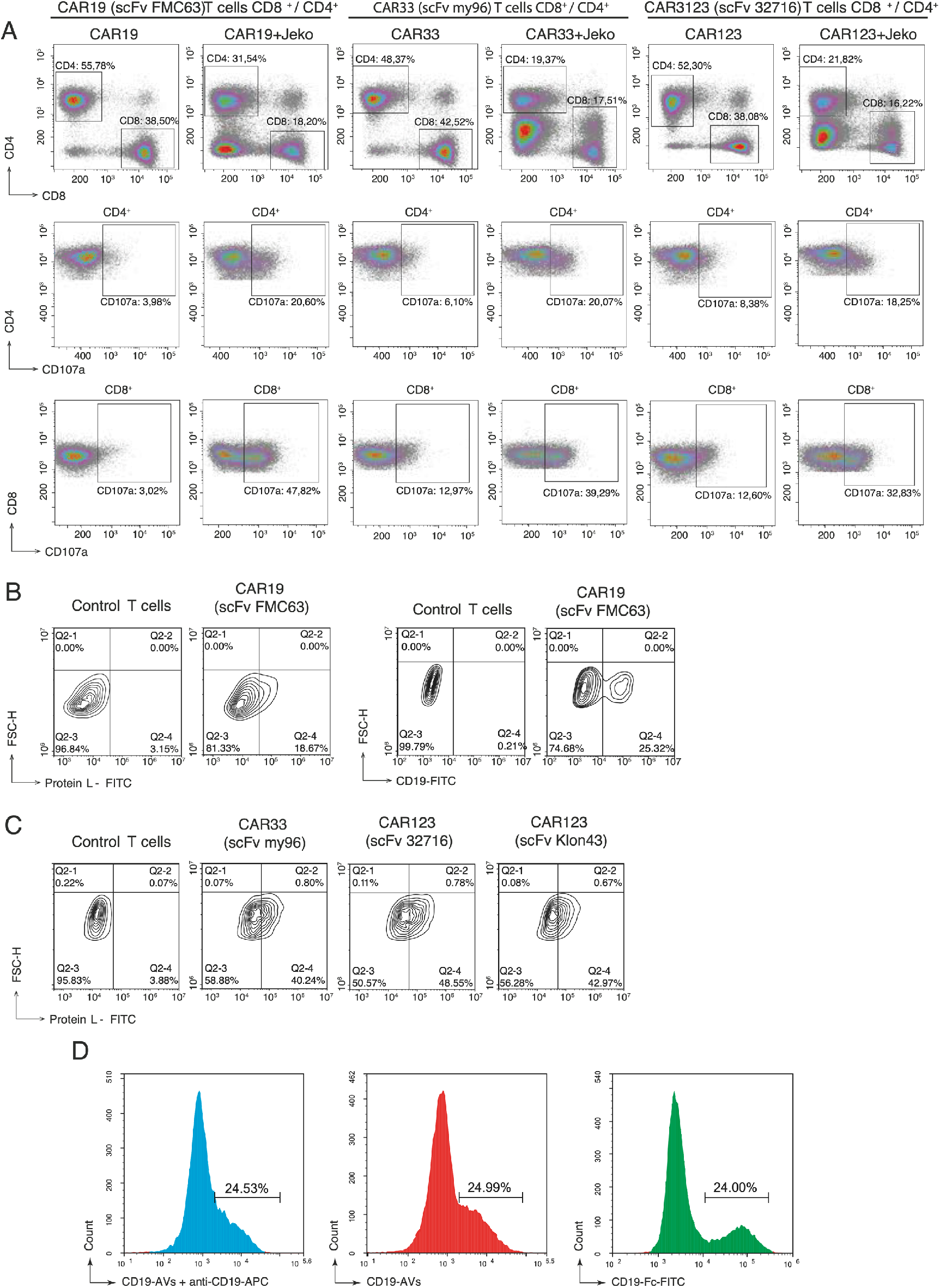
Identification of CD19, CD33, and CD123 CAR-T cells by the degranulation assay and Protein L. **(A)** CD4+ and CD8+ populations of CAR-T cells were incubated with tumor cells expressing targeted antigen and analyzed for presence of the CD107a degranulation marker. For degranulation assay 2×10^5^ of CD19, CD33, or CD123 CAR-T cells were co-incubated with Jeko-1 cells at a 1:1 ratio for 2 hours in total volume of 200 mkl. After incubation the cell mixture was washed twice (300g, 5 min) and labeled with anti-CD4, anti-CD8 and anti-CD107a antibodies for 30 min at 4°C. Subsequently, cells were washed once, analyzed by flow cytometry and plotted as two-dimensional dot plots. CAR-T cells incubated alone were utilized as a negative control. **(B)** Staining of the CD19 CAR-T cells with FITC labeled protein L and recombinant CD19-Fc protein. **(C)** Staining of the CD33 and CD123 CAR-T cells with FITC labeled recombinant protein L. **(D)** CD19 CAR-T cells were incubated with CD19-AVs and stained with anti-CD19-APC antibody.

**Supplementary Figure 4.**
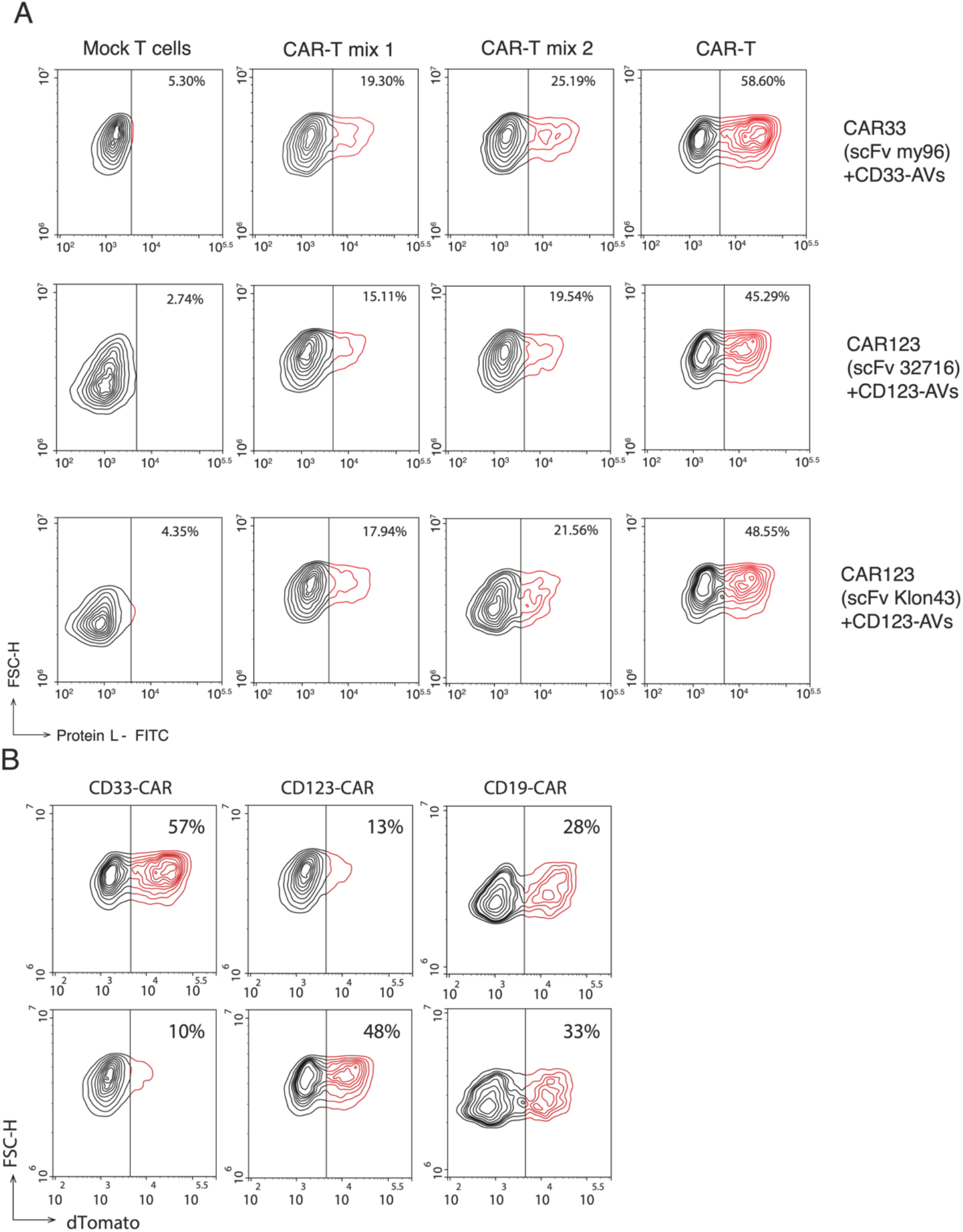
Detection of CD33 and CD123 CAR-T cells with dTomato-AVs. **(A)** Cells were stained with corresponding AVs, washed, analyzed by flow cytometry and plotted as two-dimensional contour plots. CAR demonstrates detection of (untouched) population of CAR-T cells. CAR-T mix 1 and CAR-T mix 2 represent CAR-T cells mixed with Mock T cells in a ratio of 1:2 or 1:1, respectively. Numbers indicate percentage of the detected CAR-T cells. **(B)** Double-antigen-positive HeLa/tdTomato cells can be a source of AVs with dual specificity. CD33-CD19-AVs stain both CD19 CAR-T and CD33 CAR-T cells but not CD123 CAR-T cells, while CD123-CD19-AVs stain CD19 and CD123 CAR-T cells but not CD33 CAR-T cells. CAR-T cells were stained with corresponding AVs, washed, analyzed by FC and plotted as two-dimensional contour plots.

**Supplementary Figure 5.**
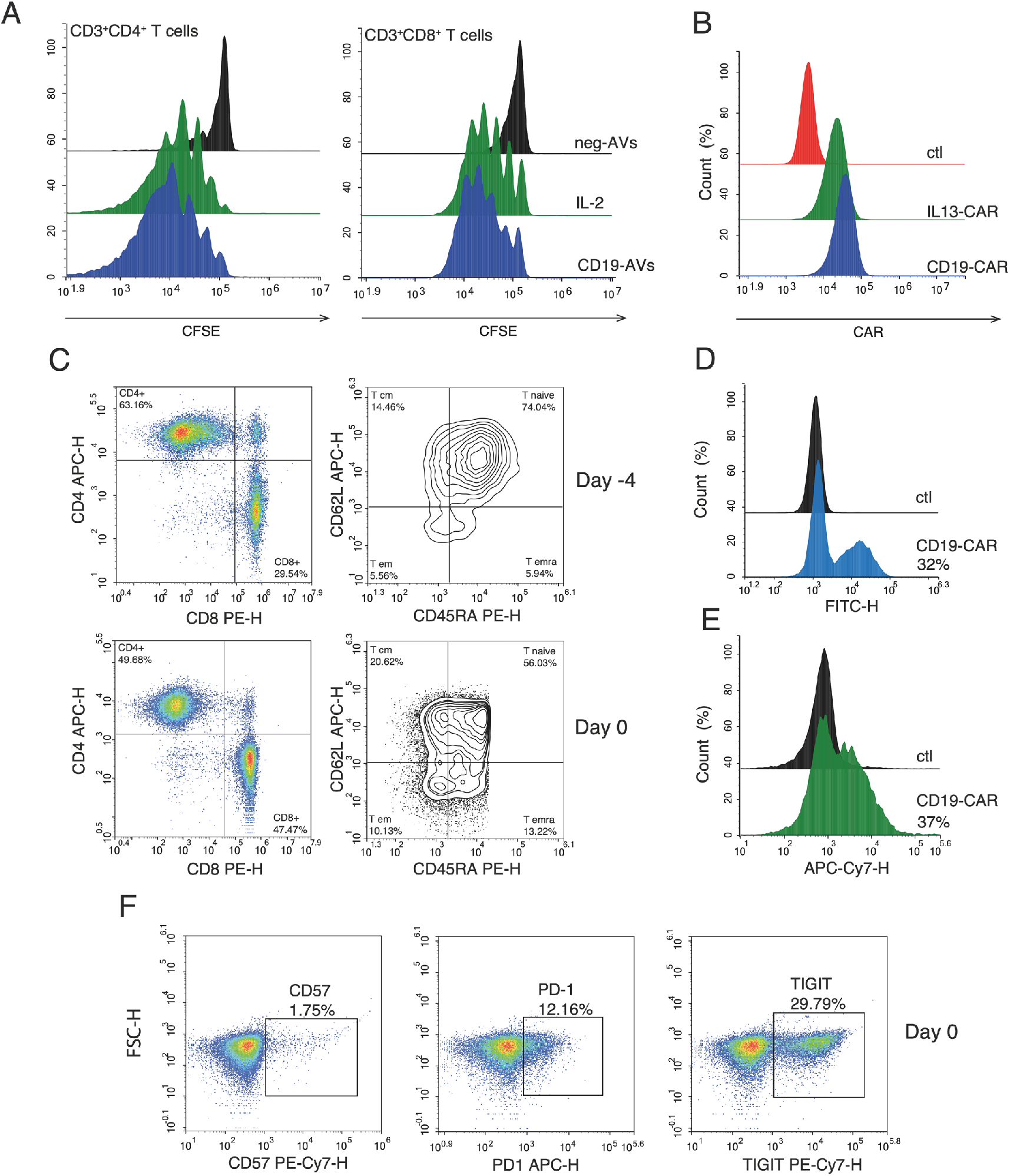
Preliminary analysis of CAR-T cells. **(A)** CFSE-based tracking of proliferation in CD4^+^ and CD8^+^ T cells that were cultured for 4 days in presence of either IL-2, or antigen-positive (CD19-AVs) or antigen-negative (neg-AVs) artificial vesicles at a ratio of 5:1. **(B)** Analysis of CD19-CAR and IL-13-CAR expression on Jurkat-NFAT-luc cells prior to CAR-T cell activation assay. **(C)** FACS quantification of changes in CD19-CAR cell surface phenotype (reflects maturation/differentiation state) and CD4/CD8 ratio on Days -4 and 0 of *in vitro* T-cell expansion. CAR-T cells were stained with corresponding antibodies, washed, analyzed by FC and plotted as two-dimensional contour plots. **(D)** Analysis of CD19-CAR expression on T cells prior to *in vitro* T-cell expansion. (E) Analysis of CD19-CAR and IL-13-CAR expression on T cells before CAR-T cell degranulation and cytotoxicity assays. (F) FACS quantification of CD19-CAR cells expressing exhaustion markers PD-1, CD57 and TIGIT among CD3^+^ cells on Day 0 of expansion experiment. CAR-T cells were stained with corresponding antibodies, washed, analyzed by FC and plotted as two-dimensional contour plots.

**Supplementary Figure 6.**
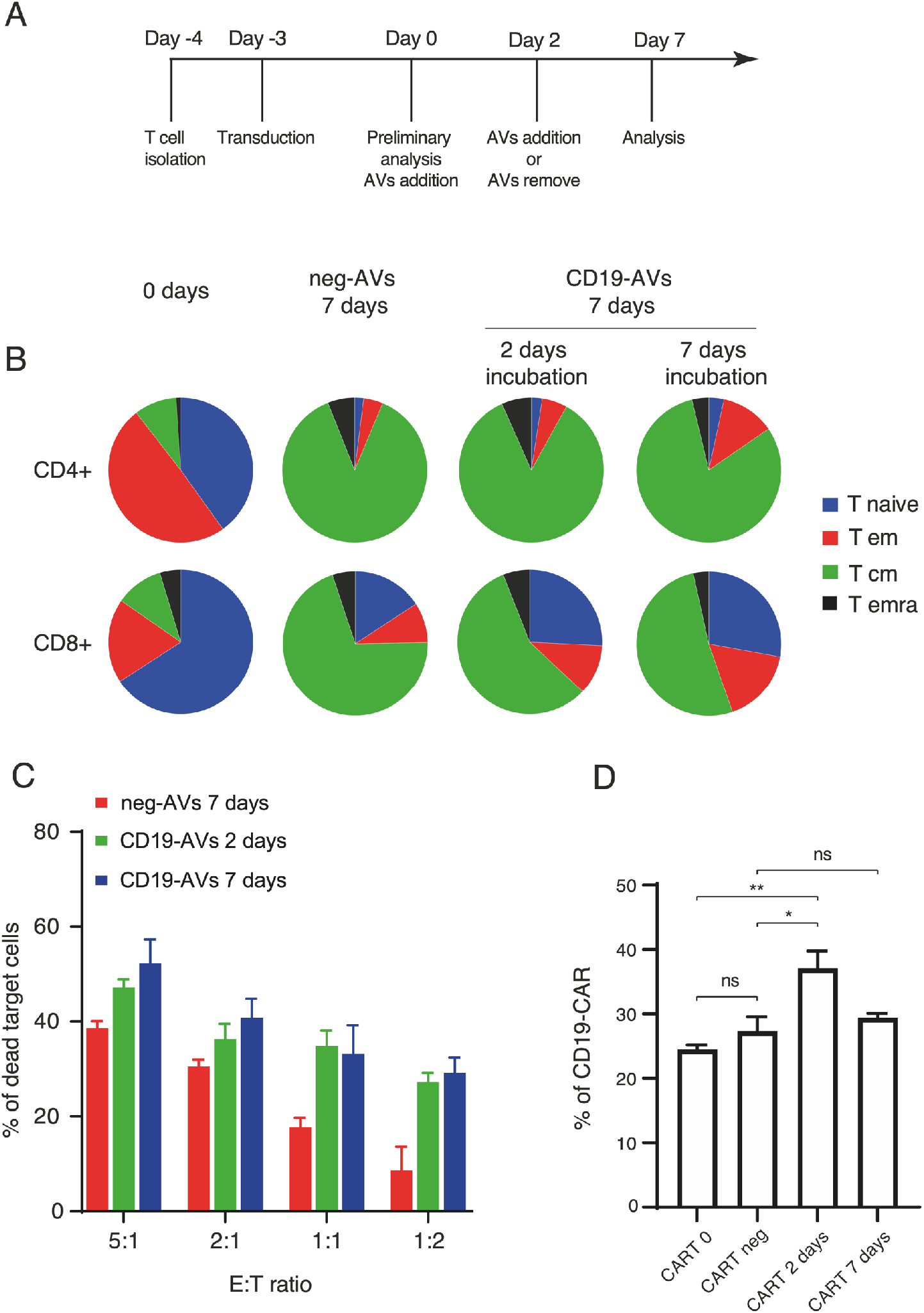
Antigen-specific expansion of CAR-T cells during generation of autologous CAR-T product. **(A)** The scheme shows major time points in the expansion experiment protocol. **(B)** FACS quantification of changes in CD4^+^ and CD8^+^ T cell surface phenotype (reflects maturation/differentiation state) in response to neg-AVs or CD19-AVs during 2 or 7 days of co-incubation at a 10:1 CAR-T:AV ratio. **(C)** *In vitro* killing activity of CD19-CAR T cells cultured with neg-AVs or CD19-AVs for 2 or 7 days at a 10:1 CAR-T:AVs ratio. At day7 after start of treatment with vesicles, CAR-T cells were incubated with Jeko-1 target cells at various effector-to-target cell ratios. Data are represented as the mean ± s.d. of three experimental replicates and are representative of at least three independent experiments. **(D)** Assesment of changes in the CAR+ subpopulation of T cells exposed to neg-AVs or CD19-AVs for 2 or 7 days at a 10:1 CAR-T:AV ratio. Data are represented as the mean ± s.d. of two experimental replicates and are representative of at least two independent experiments. Statistical analysis was performed using one-way ANOVA with multiple comparisons. For all panels, \**P* < 0.05, \*\**P* < 0.01, \*\*\**P* < 0.001, \*\*\*\**P* < 0.0001.

**Supplementary Video 1. Three-dimensional reconstructions from optical sections of CD19 CAR-T cells stained by CD19-AV using confocal microscopy**.

## References

1. Hollyman, D. et al. Manufacturing validation of biologically functional T cells targeted to CD19 antigen for autologous adoptive cell therapy. J Immunother 32, 169–180 (2009).

2. Maus, M. V. et al. Ex vivo expansion of polyclonal and antigen-specific cytotoxic T lymphocytes by artificial APCs expressing ligands for the T-cell receptor, CD28 and 4-1BB. Nat Biotechnol 20, 143–148 (2002).

3. Cheung, A. S., Zhang, D. K. Y., Koshy, S. T. & Mooney, D. J. Scaffolds that mimic antigen-presenting cells enable ex vivo expansion of primary T cells. Nature Biotechnology 36, 160– 169 (2018).

4. Zhang, D. K. Y., Cheung, A. S. & Mooney, D. J. Activation and expansion of human T cells using artificial antigen-presenting cell scaffolds. Nature Protocols 15, 773–798 (2020).

5. Xu, Y. et al. Closely related T-memory stem cells correlate with in vivo expansion of CAR.CD19-T cells and are preserved by IL-7 and IL-15. Blood 123, 3750–3759 (2014).

6. Neal, L. R. et al. The Basics of Artificial Antigen Presenting Cells in T Cell-Based Cancer Immunotherapies. J Immunol Res Ther 2, 68–79 (2017).

7. Hansen, T. H. & Bouvier, M. MHC class I antigen presentation: learning from viral evasion strategies. Nature Reviews Immunology 9, 503–513 (2009).

8. Rushworth, D. et al. Universal artificial antigen presenting cells to selectively propagate T cells expressing chimeric antigen receptor independent of specificity. J Immunother 37, 204– 213 (2014).

9. Oelke, M., Krueger, C., Giuntoli, R. L. & Schneck, J. P. Artificial antigen-presenting cells: artificial solutions for real diseases. Trends in Molecular Medicine 11, 412–420 (2005).

10. Sadelain, M., Brentjens, R. & Rivière, I. The Basic Principles of Chimeric Antigen Receptor Design. Cancer Discovery 3, 388–398 (2013).

11. Oshchepkova, A. et al. Cytochalasin-B-Inducible Nanovesicle Mimics of Natural Extracellular Vesicles That Are Capable of Nucleic Acid Transfer. Micromachines 10, 750 (2019).

12. Theodoropoulos, P. A. et al. Cytochalasin B may shorten actin filaments by a mechanism independent of barbed end capping. Biochem. Pharmacol. 47, 1875–1881 (1994).

13. Gomzikova, M. O. et al. Cytochalasin B-induced membrane vesicles convey angiogenic activity of parental cells. Oncotarget 8, 70496–70507 (2017).

14. Sun, X. et al. Surface-Engineering of Red Blood Cells as Artificial Antigen Presenting Cells Promising for Cancer Immunotherapy. Small 13, 1701864 (2017).

15. Perica, K., Kosmides, A. K. & Schneck, J. P. Linking form to function: Biophysical aspects of artificial antigen presenting cell design. Biochimica et Biophysica Acta (BBA) -Molecular Cell Research 1853, 781–790 (2015).

16. Pick, H. et al. Investigating cellular signaling reactions in single attoliter vesicles. J. Am. Chem. Soc. 127, 2908–2912 (2005).

17. Xie, J. et al. Immunochemical engineering of cell surfaces to generate virus resistance. Proc. Natl. Acad. Sci. U.S.A. 114, 4655–4660 (2017).

18. De Oliveira, S. N. et al. A CD19/Fc fusion protein for detection of anti-CD19 chimeric antigen receptors. J Transl Med 11, 23 (2013).

19. Isser, A., Livingston, N. K. & Schneck, J. P. Biomaterials to enhance antigen-specific T cell expansion for cancer immunotherapy. Biomaterials 268, 120584 (2021).

20. Wang, X. & Rivière, I. Clinical manufacturing of CAR T cells: foundation of a promising therapy. Molecular Therapy - Oncolytics 3, 16015 (2016).

21. Turtle, C. J. et al. CD19 CAR-T cells of defined CD4+:CD8+ composition in adult B cell ALL patients. J Clin Invest 126, 2123–2138 (2016).

22. Nicholson, I. C. et al. Construction and characterisation of a functional CD19 specific single chain Fv fragment for immunotherapy of B lineage leukaemia and lymphoma. Mol. Immunol. 34, 1157–1165 (1997).

23. Schneider, D. et al. A Unique Human Immunoglobulin Heavy Chain Variable Domain-Only CD33 CAR for the Treatment of Acute Myeloid Leukemia. Front Oncol 8, 539 (2018).

24. Du, X., Ho, M. & Pastan, I. New Immunotoxins Targeting CD123, a Stem Cell Antigen on Acute Myeloid Leukemia Cells. Journal of Immunotherapy 30, 607–613 (2007).

25. Albanell, J. & Baselga, J. Trastuzumab, a humanized anti-HER2 monoclonal antibody, for the treatment of breast cancer. Drugs Today (Barc) 35, 931–946 (1999).

